# Friend, food, or foe: sea anemones discharge fewer nematocysts at familiar anemonefish after delayed mucus adaptation

**DOI:** 10.1101/2024.02.22.581653

**Authors:** Cassie M. Hoepner, Emily K. Fobert, David Rudd, Oliver Petersen, Catherine A. Abbott, Karen Burke da Silva

## Abstract

For decades, it has been hypothesized that anemonefishes are able to live within the stinging tentacles of host sea anemone species because the chemical composition of their mucus layer inhibits or lacks the trigger for firing host nematocysts. However, there is very little molecular evidence for this, beyond suggestions that glycans in the mucus could be key. In this study we assessed these hypotheses by testing Bubble-tip anemone (*Entacmaea quadricolor*) nematocysts in response to three different mucus sources, before and after anemonefish association. We also profiled the corresponding mucus lipid and glycan composition of anemonefish. Host sea anemones significantly reduced nematocyst firing at acclimated anemonefish mucus compared to mucus from unacclimated individuals. Changes in anemonefish mucus glycan composition became distinguishable three weeks after introduction relative to an anemonefish that was not living in association of a host sea anemone. The glycan composition reverted back to a pre-acclimated composition when profiled 24 hours after anemonefish removal from a host sea anemone. Triggering fewer nematocysts through glycan profile alterations may be an important adaptation that has enabled anemonefish to live long-term in a sea anemone host. However, the delay in mucus response indicates it is not the initial mechanism used by anemonefish to enter a host sea anemone without being stung.

## 2. Introduction

Symbiotic relationships can take a variety of forms: mutualistic, parasitic, predatory, competitive, or commensal. Relationships between venomous and non-venomous species generally function as predator-prey, where the predator evolves more toxic venom, and in response, their prey evolve greater resistance to predatory toxins (Holding et al. 2016, Arbuckle et al. 2017); referred to as a chemical arms race. Coevolution has resulted in a huge diversity of protein and peptide toxins as well as unique strategies for prey capture and defence. The mutualism between anemonefish and sea anemones is a rare example of a venomous species and a non-venomous species both benefiting from the toxic relationship. Other mutualistic relationships do exist between venomous species and non-venomous species, such as cnidarians and zooxanthellae or bacteria (Pontasch et al. 2013, Breusing et al. 2022), but rarely does it involve a vertebrate as the counter-part to an venomous host. The mechanisms that have evolved to enable anemonefish, a potential prey species, to live within the toxic environment of host sea anemones is complex and has yet to be fully resolved, however as indicated by Mebs (2009), it likely involves the mucus layer of the anemonefish. All fish have a mucus layer that covers and protects surfaces including skin, gills and gut. The main constituent of this protective barrier is glycoproteins (Gomez et al. 2013) that are composed of more than 50% carbohydrates (Hattrup & Gendler 2008). These carbohydrates are mainly in the form of O-glycans (mucins) that protect against pathogens, and have anti-microbial functions (Reverter et al. 2018). Glycans are the sugar side chains attached to proteins that contribute to the barrier function of mucins and protect glycoproteins from cleavage by proteases (Varki 2016).

Sea anemone venom is a complex and diverse mixture of a variety of toxic components, including cytolysins (toxins that cause cell lysis), neurotoxins (toxins that damage or impair the nervous system) and phospholipases (enzymes which cause inflammation and pain) amongst others (Anderluh & Macek 2002, Frazao et al. 2012, Madio et al. 2019). Cnidarians (corals, sea anemones and jellyfish) are the only venomous organisms that do not have centralized venom glands (e.g. snakes); instead the venom is produced in tissues throughout their body via nematocytes and ectodermal gland cells (Madio et al. 2019). Compared to other venomous animals, relatively little is known about cnidarian venom production or delivery. Sea anemones utilise cells (cnidae), containing highly specialised and complex stinging organelles (cnidocytes) to distribute venom and these are deployed to capture prey (typically fish and arthropods) and for defence purposes. The class Anthozoa (which comprises all sea anemones) (Anderson & Bouchard 2009) has three types of cnidocytes: nematocyst, spirocysts, and ptychocysts. Studies have shown that sea anemones can have multiple nematocyst types (six types, two of them exclusive to the class) that are found throughout various tissues (tentacle, filament, column, actinopharynx) (Krayesky et al. 2010, Jindrich 2011) that disperse the sea anemones’ venom. The tentacles of sea anemones have been found to only exhibit spirocysts and two types of nematocysts: microbasic p-magistophore and basitrichous (Fautin 1981, Krayesky et al. 2010, Jindrich 2011).

The discharge of nematocysts are controlled by chemosensory, mechanosensory and neurological pathways that respond to external sensory stimulation (Anderson & Bouchard 2009). Sea anemones possess chemoreceptors that can detect N-acetylated sugars present in the mucus layer of fish (Abdullah & Saad 2015). When N-acetylated sugars, specifically the acidic side chain of the glycoprotein, binds to the sea anemone’s chemoreceptor, a multiple signal pathway is triggered resulting in the firing of nematocysts (Anderson & Bouchard 2009). It was previously thought that the firing of nematocysts was not controlled by the sea anemone, but Conklin and Mariscal (1976) found that sea anemones can adjust the number of nematocysts fired under experimental conditions. Conklin & Mariscal (1976) also found that sea anemones will reduce the number of nematocysts fired when satiated (Conklin & Mariscal 1976). The synthesis of toxins and nematocysts have a metabolic cost (Fautin 2009, Sachkova et al. 2020, Kaposi et al. 2022), resulting in optimization of nematocyst use toward different predators and prey in different ecological contexts.

While sea anemones utilise their nematocysts to capture prey and defend themselves from predators, their symbiont, the anemonefish, is able to live unharmed amongst the sea anemone’s venomous tentacles for the duration of the fish’s post-larval life (Mebs 1994, 2009). There are approximately 1170 sea anemone species (Rodríguez et al. 2022), yet only ten species from three families (*Thalassianthiade, Actinidae, Stichodactylidae*) form associations (as hosts) with one or more of the 28 species of anemonefish (*Amphiprion*) (Fautin 1991, Burke da Silva & Nedosyko 2016, Tang et al. 2021). The sea anemone provides a safe site for anemonefish reproduction and protection from predation, whereas the anemonefish helps to increase the growth, reproduction, and defense of host sea anemones by providing nutrients from their feces and increased oxygenation by swimming amongst the sea anemones tentacles and chasing off potential sea anemone predators (Szczebak et al. 2013, Frisch et al. 2016, Schligler et al. 2022). As anemonefish contribute to predator defence of host sea anemones, we hypothesise that host sea anemones will reduce their reliance on chemical defence when in association with an anemonefish.

Much debate has occurred regarding the mechanisms that enable anemonefish to survive in a venomous sea anemone host (Fautin 1991, Mebs 2009, Burke da Silva & Nedosyko 2016), but to date the exact mechanism remains unknown. One of the key hypotheses suggests that anemonefish mucus either inhibits or lacks the trigger for the firing of sea anemone nematocysts (Lubbock 1980, Abdullah & Saad 2015). Lubbock (1980) qualitatively observed the behavioural response of the host sea anemone, Haddon’s anemone (*Stichodactyla haddoni*), when presented with mucus from different anemonefish and damselfish species. When Clarke’s anemonefish (*Amphiprion clarkii*) was both associated with Haddon’s anemone, and not associated, the anemonefish mucus (presented on a glass rod) did not elicit a strong behavioural response from the sea anemone (0/10 and 8/45, respectively). In comparison, mucus from closely related damselfish (*Chromis caerulea, Dascyllus aruanus, Paraglyphidodon nigroris*), always elicited a strong behavioural result (35/35). A more recent study by Abdullah and Saad (2015) found that mucus from *A. ocellaris* contained significantly lower concentrations of N-acetylated sugars (Neu5Ac) compared to the mucus of other coral reef fish (*Abudefduf sexfasciatus* and *Thalassoma lunare*). Neu5Ac is a sialic acid side chain that has been shown to trigger the chemoreceptors that control sea anemone nematocyst firing (Ozacmak et al. 2001, Abdullah & Saad 2015). However, no study to date has quantified sea anemone nematocyst response to anemonefish mucus (as opposed to the qualitative work by Lubbock (1980)).

In this study, we investigate the role sea anemone nematocysts play in facilitating the symbiotic relationship with anemonefish and the lipid and glycan content of anemonefish mucus. Specifically, we build on the research by Abdullah and Saad (2015) by testing the hypothesis that anemonefish mucus lacks the trigger for stimulating nematocyst firing, thus facilitating anemonefish-sea anemone symbiosis. To test this hypothesis, we use a manipulative lab experiment to introduce mucus collected from 1) a symbiotic anemonefish (*Amphiprion percula*), 2) a non-symbiotic damselfish (*Chromis viridis*), and 3) a sea anemone food source (prawn), to investigate the impact on nematocyst firing. We also test *Entacmaea quadricolor* nematocyst response to all three mucus sources before and after anemonefish association, to determine if anemonefish acclimation alters sea anemone nematocyst response. Further, we expanded on research by Heim et al. (2023) to examine the lipid and glycan profile of anemonefish mucus before and after association with a host sea anemone via metabolomic profiling.

## 3. Materials & Methods

### 3.1 Study species and experimental set-up

Twelve pairs of anemonefish (n=24) *(Amphiprion percula)* were purchased from Cairns Marine and six *E. quadricolor* sea anemones, obtained from a local aquarium store in Adelaide (harvested in Cairns), were transported to the animal house facility at Flinders University, South Australia in 2019 for the metabolite experiment. In 2020, ten host sea anemones (*Entacmaea quadricolor*) (∼5-7cm diameter) were obtained from an aquarium store in Adelaide South Australia (harvested from Western Australia) and twelve Blue Green Chromis damselfish (*C. viridis*), also from an aquarium store in Adelaide South Australia for the nematocyte experiment. Ten pairs of *A. percula* anemonefish (female and male) (n=20) were reused from the metabolite experiment. For acclimation to the animal house upon arrival, sea anemones were held in individual 30L tanks for a 2-week acclimation period (26.5 °C ± 0.7, salinity 37.5 ± 1.5, pH 7.91 ± 0.2). Each sea anemone was fed a small piece of prawn every three to four days except in the 48 hours leading up to each sampling event. Each tank had a Fluval Aquatic Marine Nano 3.0 lights (2500 lux on a 12:12 L:D light cycle). Anemonefish were housed in 30L tanks that contained a terracotta pot (a sea anemone surrogate) for a 2-week acclimation period prior to the experiment. Recirculating tanks (30L) holding anemonefish pairs were attached to a sump system separate to the sea anemones (27 °C ± 0.6, salinity 36.5 ± 1.5, pH 8.01 ± 0.2). Damselfish were held in an isolated 200L holding tank for a 2-week acclimation period, then separated into individual 30L recirculating tanks attached to a separate sump system to the anemonefish and sea anemones, for the nematocyte experimental period. *A. percula* and *C. viridis* were fed twice daily with commercial pellets (Hikari Marine S) and mysid shrimp. For the metabolite experiment six pairs of *A. percula* were randomly assigned to either the control or treatment groups. Control and treatment tanks were on a separate, recirculating water system (control: 27 °C ± 0.6, salinity 36.5 ± 1.5, pH 8.01 ± 0.2, treatment: 27 °C ± 0.6, salinity 36.5 ± 1.5, pH 8.01 ± 0.2).

### 3.2 Sample preparation

Glass microscope slides were used to collect mucus samples. For the nematocyte experiment each glass slide was marked with a glass pen at 2.5cm from the bottom of the slide to create a 2.5 cm^2^ area for sampling (following methodology used by Pantin 1942, Conklin & Mariscal 1976, Mauch 1998, Greenwood et al. 2004, Todaro & Watson 2012). Glass slides were prepared prior to sampling by dipping the slide into molecular grade 100% ethanol (Sigma-Aldrich 200-578-6) for 10 seconds for sterilisation and placed into a 50mL falcon tube to air dry. *A. percula* and *C. viridis* were individually collected in a hand-net, and mucus was collected by gently scraping the sampling area of the glass slide along the side of the fish from operculum to tail; one glass slide was used for each side of the fish (approved by the Flinders University Animal Ethics Committee E470-18). Light scraping was used to avoid the collection of epidermal skin cells. For the prawn mucus, a small fleshy piece of defrosted prawn was rubbed onto the slide sampling area. All slides were left to dry to prevent mucus washing from the slide. Imaging from SEM clearly shows full mucus cover on all three slide types, providing a 2-dimensional surface to introduce to the anemone (Supp Fig 1). We tested our method for introducing a slide to sea anemones by placing a blank slide, no mucus (n=10) within the sea anemone tentacles. We did not include blank controls at each of the timepoints because zero nematocysts were triggered by the sea anemones in all ten replicates.

For the metabolite experiment the full glass slide was used with only *A. percula* anemonefish sampled. Both sides of the fish were scraped using the same slide to obtain the most mucus possible for the sample and the slides were then transferred into a 50mL tube and immediately frozen at −80°C until processing.

### 3.3 Nematocyte experiment

#### 3.3.1 Sampling design

Nematocyte response was measured across a nine-week period (Fig 1). In week 1, each sea anemone (n=10) was randomly assigned to one of the three mucus treatment types: *anemonefish*, *damselfish*, and prawn. Two slides were introduced to each sea anemone, one at a time, to induce nematocyst firing (Conklin & Mariscal 1976, Hoepner et al. 2019). The slides were placed mucus side facing down into the tentacles and held for a period of five seconds on the opposite side of the sea anemone to ensure new tentacles were being exposed to the stimulus. This process was repeated in weeks 2 and 3, alternating the treatment slide introduced so that each sea anemone was tested against all three treatments across the three weeks. Following the ‘before’ association period, a pair of *A. percula* (female and male) were introduced into each tank with a sea anemone and left for three weeks to establish symbiosis. All anemonefish became established within sea anemones within 48 hours after introduction and remained living within the tentacles for the 3-week period, illustrating full association. The three treatment types were repeated over a further three weeks, at week 7, 8 and 9 for the ‘after’ association period. In this study hereafter, references to the before and after period of the BACI design will be referred to as ‘non-hosting’ and ‘hosting’ periods to reflect the nature of symbiosis at this time. The anemonefish mucus used to elicit nematocyst firing was collected only from the female (larger) anemonefish (n=10) living in the same tank with a sea anemone host, thus the sea anemones were responding to ‘familiar’ anemonefish mucus. The same anemonefish was also presented to the same sea anemone in both the non-hosting and hosting periods.

**Figure 1:**
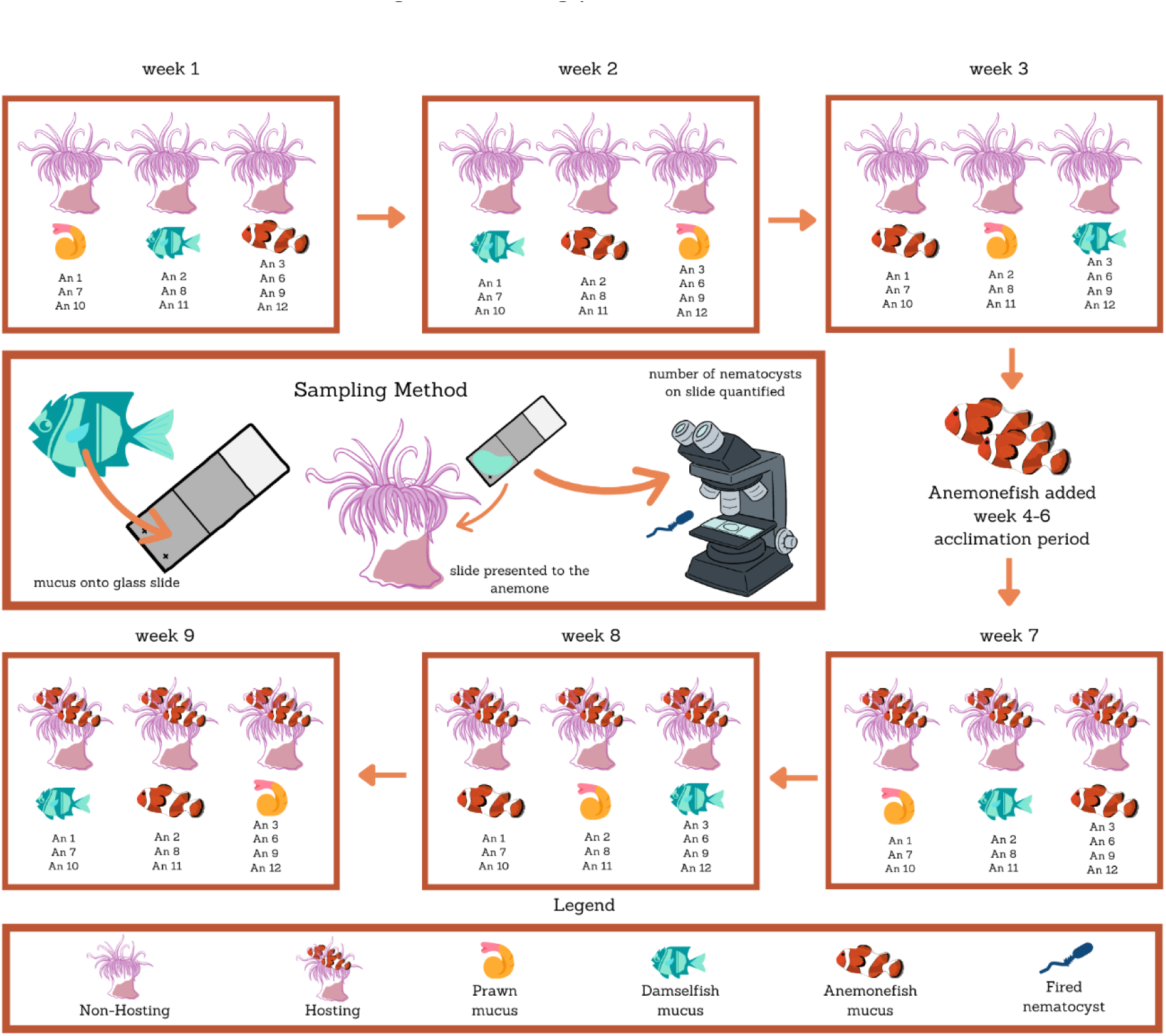
Experimental design showing mucus types introduced to *Entacmaea quadricolor* anemones (n=10) across the nine-week experimental period to assess nematocyte discharge during hosting and non-hosting periods. Each mucus type (anemonefish, damselfish, prawn) was presented to each anemone over the three-week non-hosting period. A pair of anemonefish were then added for a three week acclimation period. Mucus sampling was then repeated over three weeks for the hosting period, with ‘familiar’ anemonefish mucus introduced to the corresponding anemone.

#### 3.3.2 Nematocyte Counts

After a slide was exposed to a sea anemone, it was stained using methylene blue (0.5g in 100mL H_2_O) and allowed to air-dry. Fired nematocysts capsules were counted under a Zeiss compound microscope at 400x magnification. All nematocysts were counted within the marked 2.5 cm^2^ area of the glass slide by moving at 1mm intervals back and forth across the slide. As two slides for each treatment were introduced to a sea anemone at each time interval, the number of capsules fired was calculated as the mean number of nematocysts across the two slides. Sea anemone behavior was observed during the slide introduction, and in instances where the sea anemone retracted its tentacles during sampling or remained contracted during the sampling day (*A. percula* Before =1, *C. viridis* Before =1, Prawn With =2), data was excluded as this did not represent a typical predator/defence response. In all instances, slides that were excluded based on sea anemone behavior contained <10 nematocysts. When examining nematocysts, we defined ‘fired’ as a nematocyst that has been fired from the tentacle of the sea anemone, ‘unfired’ as a nematocyst that was still intact within the sea anemone tentacle, ‘discharged’ as a nematocyst that had been fired and that had expelled its internal contents (thread etc.) and ‘undischarged’ as a nematocyst that has been fired but has not expelled its internal contents (Supp Fig 2).

### 3.4 Metabolite experiment

#### 3.4.1 Sampling design

Mucus from *A. percula* was sampled weekly, for a period of 8 weeks from both fish in association with sea anemones (hosted treatment) and fish without a sea anemone (non-hosted control) (approved by the Flinders University Animal Ethics Committee E470-18) (Fig 2). Mucus from all 12 pairs was sampled at week 0 for an initial mucus sample. One week later an *E. quadricolor* anemone with a tile for attachment was added to each of the six treatment tanks and the terracotta pot was removed from these tanks. Anemonefish mucus was sampled again for both the treatment and control groups 48 hours after sea anemones were added to the tank and again after 1, 2 and 3 weeks. One-week later all sea anemones were removed from the treatment groups and the terracotta pots re-added. Mucus was re-sampled for both the control and treatment groups 24 hours followed by 1, 2 and 3 weeks after sea anemone removal. Throughout the entire 8-week experiment anemonefish in the control group were kept without an anemone and with a surrogate terracotta pot.

**Figure 2:**
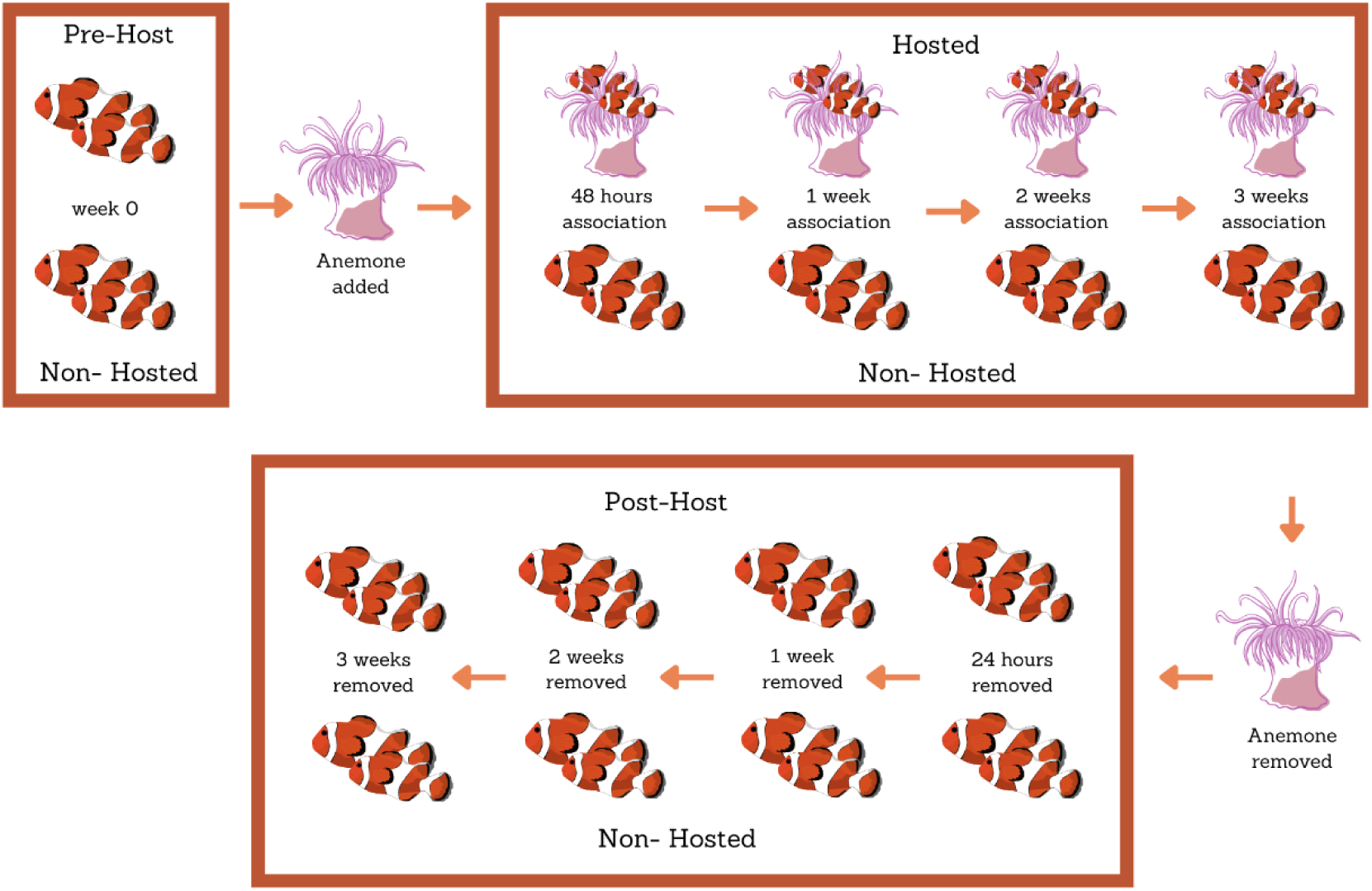
Mucus collecting procedure for *Amphiprion percula* anemonefish with and without *Entacmaea quadricolor* sea anemone presence over the 8-week experimental period, for hosted (n=12) and non-hosted (n=12) groups.

#### 3.4.2 MALDI TOF/TOF analysis

The samples underwent metabolomic analysis at the Melbourne Centre for Nanofabrication, Monash University. To prepare the samples for analysis a chloroform/methanol phase partition was performed separating each anemone fish mucus sample into a lipid and aqueous phase, which was stored at −80°C before use. The aqueous phase contained soluble proteins largely glycoproteins and was kept for glycan analysis.

For the lipid phase, matrix-assisted laser desorption/ionization time-of-flight (MALDI-TOF/TOF) mass spectrometry (MS) (Bruker Germany) measurements were performed to identify changing molecular ions. 2x 4.5µl of each sample was pipetted onto a novel nanofabricated surface (Minhas et al. 2020) for MALDI analysis. A total of 5000 hits was used to create a spectrum for each sample. Using FlexAnalysis (Bruker, Germany) each spectrum was background subtracted and had the signal-to-noise ratio set to 7 (centroid peak picking). Samples were then peak binned using MetaboAnalyst (www.metaboanalyst.ca) (Pang et al. 2020) with a mass tolerance of 1 m/z. Samples were median normalised, log-transformed and centre-scaled.

For the aqueous phase containing soluble glycoproteins, samples were thawed and pooled (n=3-4 per pool) into nine duplicate timepoint groups (n=18) for each of the treatment groups (Wk 0; 48h, Wk 1, 2 and 3 association with host; 24 hrs, 1, 2 and 3 weeks removed from host) and pooled into three groups for the control group (n=7-11). Amicon® Ultra 0.5mL 3K spin columns (Merck KGaA, Darmstadt, Germany) were used to concentrate the pooled samples before N-glycan digestion. The pooled samples were loaded onto the Amicon® spin column and spun at 14,000 g for 20 mins. All proteins/peptides < 30 aa were discarded in the flow through. Concentrated proteins >3000 Da were washed with 400 µl UltraPure^TM^ DNase/RNAse-Free Distilled Water (Invitrogen^TM^, 10977015) and spun for 14,000g for 25 mins. Approximately 40 µl of retentate was recovered for each sample.

Peptidase N-Gylcosidase F (PGNase F) was used to cleave the innermost Glc NAc and asparagine residues of the mannose, hybrid and complex oligosaccharides from the N-linked glycoproteins in our sample. Approximately 35 µl of the concentrated pooled sample was added to 10 µl of 1 x Phosphate Buffered Saline (PBS) pH 7.4 (Gibco^TM^,70011044), 50 µl of UltraPure^TM^ water and 250 units recombinant PGNase F (Glycerol-free) (New England Biolabs, Ipswich, MA, USA) in a spin column and incubated overnight in a beadbath at 37°C. Following glycan digestion, the Amicon® Ultra 0.5mL 3-K spin columns were utilised again this time to remove the PGNase F and the proteins that had been removed of glycans, and the glycans assumed to be < 3K were collected in the flow-through following centrifugation at 14,000g for 30 mins. The samples were freeze-dried down to 5µl to further concentrate samples for mass spectrometry. Two µl of each sample was mixed with 2µl of matrix (10mg/ml 2,5 dihydroxybenzoic acid (Fluka Analytical), in 50% Acetonitrile, 50% H_2_O) and 1ul was pipetted onto FlexiMass-DS disposable targets for matrix-assisted laser desorption/ionization time-of-flight (MALDI-TOF/TOF) mass spectrometry (MS) (MALDI-7090, Shimadzu). A total of 5000 hits was used to create a spectrum for each pooled glycan sample. Using MMass (Strohalm et al. 2008) each spectrum was background subtracted and spectrum smoothed. Samples were then peak binned using MetaboAnalyst (www.metaboanalyst.ca) (Pang et al. 2020) with a mass tolerance of 1 m/z. Samples were median normalised, log-transformed and centre-scaled. Accuracy ±0.05da

### 3.5 Data Analysis

All statistical analyses were undertaken using the statistical software R using base R functions (R Core Development R Core Development Team 2013, R Core Development Team 2013), except where otherwise stated. To assess how nematocyte firing differed between treatments, we used a linear mixed effects (LME) model using the ‘lmer’ function in the R package ‘lme4’ (Bates et al. 2014) with mean capsules as the response variable, and mucus treatment (anemonefish, damselfish, prawn) and experimental period (before association, with association) included as fixed factors in the model. Additionally, individual sea anemone ID was included as a random factor to account for non-independence among repeat observations, and sampling week was included as a random factor to account for any effects of time on nematocyte firing. Significant differences between pairs of group means were determined using Bonferroni post hoc analysis. Distance matrices were generated using vegdist in the R package “vegan” with a bray dissimilarity index (Dixon 2003). Principal Coordinate Analysis (PCoA) was performed using the R package “ape” (http://ape-package.ird.fr/). POST HOC PERMANOVA was performed to test for significance using pairwise.adonis2 and SIMPER was used to identify ions driving the dissimilarity using simper both from the “vegan” R package. Graphs were created in R used the package “ggplot2” (Wickham 2016). Line graph of average sequence profile of glycans was created in excel.

## 4. Results

### 4.1 Quantitative analysis of nematocyte firing in response various stimuli

The interaction term between mucus treatment and experimental period was significant (x^2^= 7.96; df =2, p <0.0187) (Fig 3, Table 1), indicating that the number of nematocyst capsules fired by *E. quadricolor* was dependent on both stimuli presented and whether the sea anemone was hosting an anemonefish. Specifically, the experimental period affect indicates that the mean number of nematocyst capsules fired was significantly lower in response to anemonefish mucus when in association with a sea anemone (mean diff =510.3, df =22.3, p<0.0185) compared to the mucus of anemonefish that were not in association with a host sea anemone. No significant differences were found in nematocyst firing at mucus from *C. viridis* and prawn treatments before and during the association periods. In the before association period, both *A. percula* and *C. vidris* mucus stimulated high nematocysts firing, (820.5 ± 470.8 nematocysts and 1151.7 ± 385.4 respectively). The food source – prawn mucus, triggered a lower number of nematocysts (275.3 ± 370.4) compared to the fish stimuli in both time periods. However, when the anemonefish were in association with their sea anemone host, *A. percula* mucus triggered significantly fewer nematocysts (320.6 ± 253.9), whilst both the non-symbiont fish *C. vidris* and the prawn mucus triggered roughly the same number of nematocysts as during the before association period (897.5 ± 258.8 and 369.1 ± 402.1 respectively).

**Figure 3:**
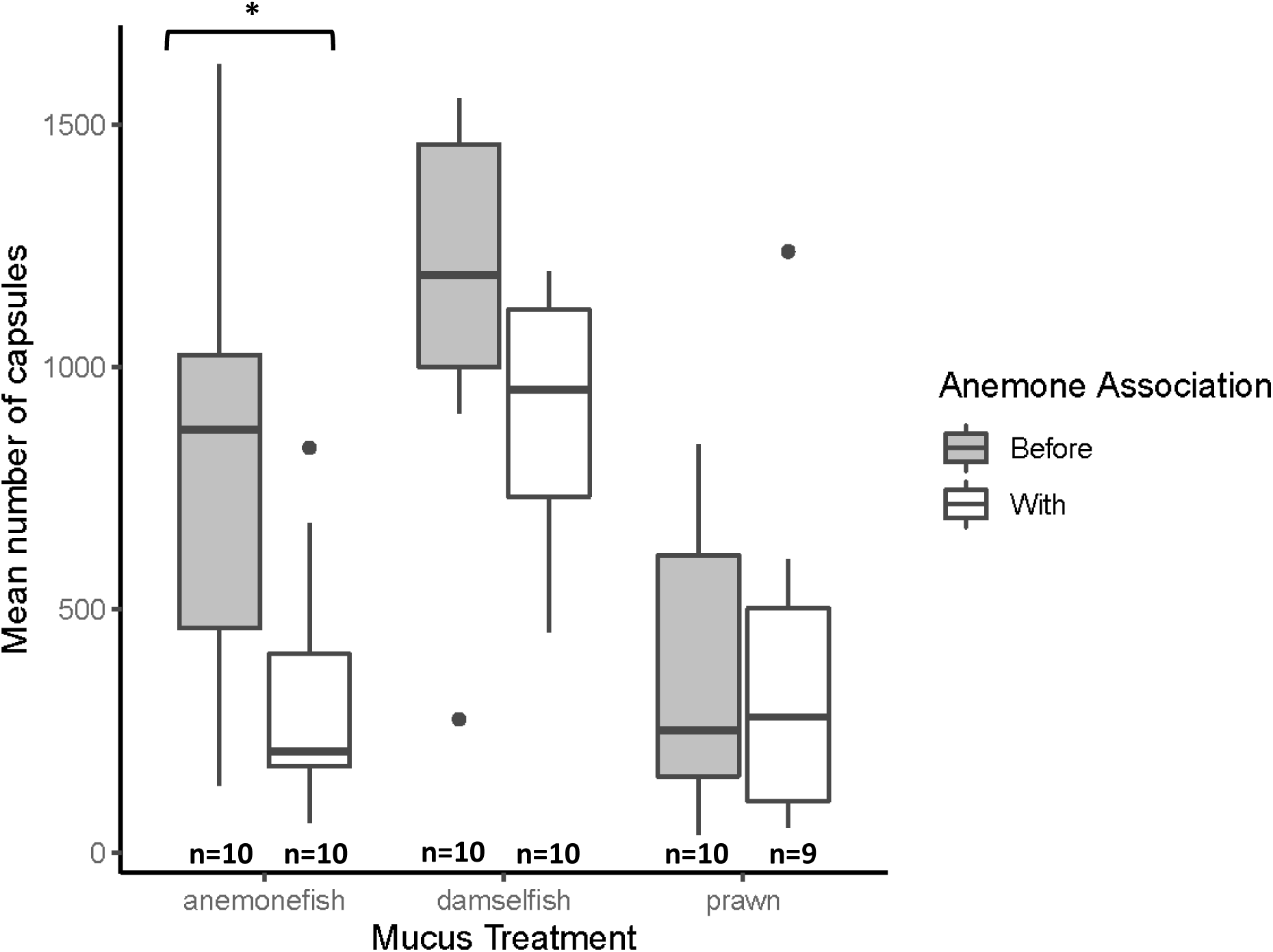
Mean number of nematocyte capsules fired by *Entacmaea quadricolor* during experimental periods (hosting and non-hosting), and mucus treatments (symbiotic *Amphiprion percula* (anemonefish), non-symbiotic *Chromis viridis* (damselfish), food source (prawn)). Boxplot shows the median and interquartile ranges. Significant differences between mean number of capsules are shown by an asterix (*). A single asterix (*) represents a significance level of p<0.01. Dots (·) represent outliers.

**Table 1:** Output from linear mixed effects model testing for differences in *Entacmaea quadricolor* nematocyte response between experimental periods (hosting and non-hosting), and mucus treatments (*A. percula*, *C. viridis*, prawn). P-values derived from Wald chi-square test with type II sums of squares. Significance is indicated in **bold**.

| **Random effects** |  |  |  |
| --- | --- | --- | --- |
| *Term* | *Variance* | *SD* |  |
| Anemone ID | 0 | 0 |  |
| Week (intercept) | 18625 | 136.5 |  |
| Residual | 87898 | 296.5 |  |
| **Fixed effects** |  |  |  |
| *Term* | $\chi^2$ | *df* | *P* |
| **Mucus treatment** | **46.44** | **2** | **8.255e-11** |
| Experimental period | 3.23 | 1 | 0.07248 |
| **Mucus treatment X experimental period** | **7.96** | **2** | **0.01872** |
| **Post Hoc – Bonferroni** |  |  |  |
| *Term* | *Mean diff* | *df* | *P* |
| ***A. percula* non-hosting/hosting** | **510.3** | **22.3** | **0.0185** |
| *C. viridis* non-hosting/hosting | 233.3 | 22.3 | 0.2574 |
| Prawn non-hosting/hosting | -26.9 | 23.5 | 0.8961 |
| Non-hosting *A. percula* / *C. viridis* | -294.9 | 47.1 | 0.0950 |
| **Non-hosting *A. percula* /prawn** | **461** | **47.1** | **0.0048** |
| **Non-hosting *C. viridis* /prawn** | **756** | **47.1** | **<0.0001** |
| **Hosting *A. percula* /*C. viridis*** | **-571.9** | **47.1** | **0.0004** |
| Hosting *A. percula* /prawn | -76.1 | 47.7 | 0.8545 |
| **Hosting *C. viridis* /prawn** | **495.8** | **47.7** | **0.0030** |

### 4.2 MALDI TOF/TOF analysis of anemonefish mucus profile in response to host sea anemone association

Using MALDI-TOF-MS we detected 65 unique lipid features (mass between 212.275 and 701.6635 Da) in *A. percula* mucus. Association with a host sea anemone did not affect the overall lipid composition of anemonefish mucus, as there was no significant difference between the lipid profiles before and during association with a host sea anemone (Supp Fig 3).

Using MALDI TOF-MS we detected 37 unique glycan features (mass between 437.41 and 1373.39 Da) across the *A. percula* mucus samples. A Principal Coordinate Analysis (PCoA). showed that association with the sea anemone *E. quadricolor* significantly altered the glycan composition of *A. percula* mucus (Fig 4A). After three weeks of association with *E. quadricolor* the glycan profile of *A. percula* mucus sits apart from all other samples (p<0.043) (Fig 4B). This is the only time point that shows change during association with *E. quadricolor* (Fig 4C), as after removal from the sea anemone resulted in the glycan profile reverting back to its original state in 24 hours (p<0.012) (Fig 4D). There were seven glycans whose percentage dissimilarity drove the significance observed (Fig 4F). Three of these glycans increased during association with a sea anemone and four decreased.

**Figure 4:**
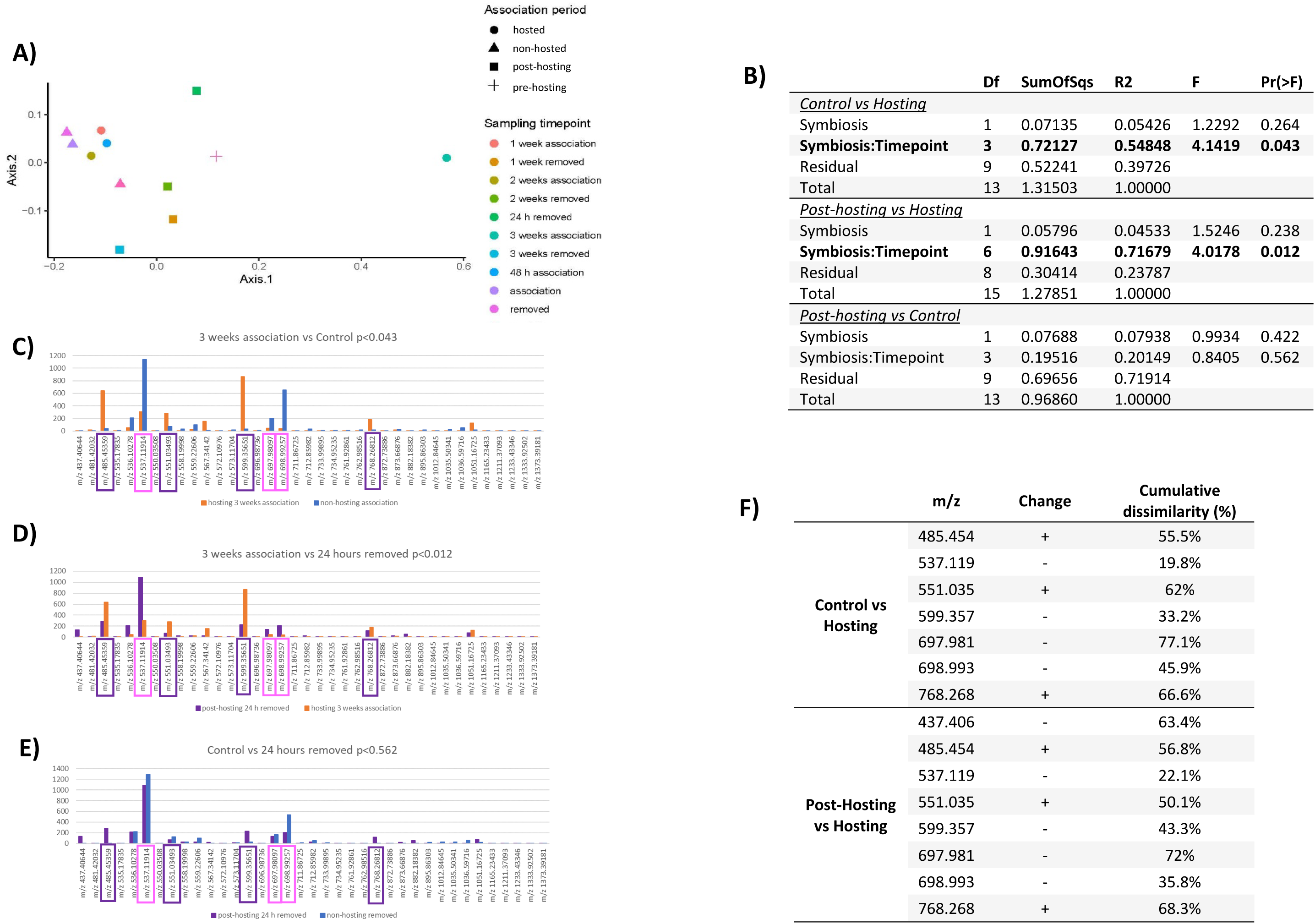
A) PCoA of *Amphiprion percula* mucus glycan profile across hosted and non-hosted periods with *E. quadricolor.* B) Post Hoc PERMANOVA of glycan composition between association periods and timepoints (**Bold** indicates significance). C-F glycans significantly altered after 3 weeks of *E. quadricolor* association + increase when hosted, - decrease when hosted with dissimilarity percentage from SIMPER, compared to control and 24 hours removed samples.

## 5. Discussion

For decades, it has been hypothesized that anemonefish are able to live within the stinging tentacles of host sea anemone species because the chemical composition of their mucus layer inhibits or lacks the trigger for the firing of sea anemone nematocysts (Lubbock 1980, Abdullah & Saad 2015). This study provides the first empirical evidence that it is only the mucus of anemonefish that are acclimated to and living in association with a host sea anemone that elicits a reduction in the number of nematocytes fired at them. In contrast, anemonefish that were not in association with a host sea anemone are very susceptible to nematocyst firing. This study also provides the first empirical evidence that the glycan profile of anemonefish is significantly altered after three weeks of living in association with a host sea anemone as proposed by Abdullah and Saad (2015), & Marcionetti et al. (2019). However, 24 hours after removal from a host sea anemone r, the glycan profile of anemonefish reverts back to what was observed when the anemonefish was not in association with the sea anemone. The delayed adaptation and quick switch-off after removal indicates that changes in the glycan profile are not the primary mechanism that anemonefish use to enter their host sea anemone without being stung, but rather is a subsequent adaptation facilitated after three weeks of living in association.

Prior to association (non-hosting) host sea anemone nematocyst response to anemonefish stimuli was similar to the response of a close non-symbiotic relative of anemonefishes, the blue green damselfish (*C. viridis*), where mucus from both species elicited a high level of nematocyst firing. However, after a three-week acclimation period with a host sea anemone, where the anemonefish performs a range of behaviours that enable them to enter, and live within the host sea anemone (Balamurugan et al. 2015), the number of nematocysts fired at the mucus of an associated anemonefish decreased significantly, while nematocyst firing remained high in response to the mucus from the non-symbiotic fish species *C. viridis*. This finding disputes results from Lubbock (1980), who found no significant difference between the observed behavioural (qualitative) response of the host sea anemone *S. haddoni* when presented with acclimated (familiar) and unacclimated mucus from the anemonefish species *A. clarkii*. Our study indicates that qualitative observation cannot accurately predict nematocyst response to stimuli and that quantitative methods are required, as we did not visually detect any difference in the physical response of the host sea anemone to different stimuli or different hosting states.

Heim et al. (2023) compared the lipid profile of anemonefishes’ and damselfishes’ mucus and found that the sphingolipid class of ceramides was a specific feature of anemonefishes’ lipid mucus composition. Heim et al. (2023) suggested monitoring changes in ceramide content of anemonefishes’ mucus when in association with a host sea anemone to determine whether anemonefishes’ lipid content is affected by association. Lipid composition across mucus samples varied considerably with no specific lipid ions showing a clear change in response to association, a pattern not identifiable as suggested by Heim et al. (2023). Determining what is endogenously produced by the anemonefish and contributions from the external environment would also be difficult to define experimentally; the changing anemonefish transcriptome may be a more reliable measure of upregulation of specific lipid metabolites. However, we did find a significant change in the glycan profile of *A. percula* mucus after three weeks association with a host sea anemone. Seven key glycans in the anemonefish mucus changed expression, three increased and four decreased when in association with a sea anemone. There is a distinct lack of data on the metabolome of marine fishes (Reverter et al. 2017, Reverter et al. 2018, Heim et al. 2023), with even less known about glycan composition, thus we were unable to identify the types of glycans represented by these altered glycoproteins due to the low quantities found in mucus.

Previous studies have indicated that the glycan content of anemonefish mucus may be involved in the reduction seen in nematocyst firing. Abdullah and Saad (2015) analysed the chemical composition of anemonefish (*A. ocellaris*) mucus and found that it contained significantly lower concentrations of Neu5Ac compared to the mucus of other coral reef fish species (*Abudefduf sexfasciatus* and *Thalassoma lunare*). Neu5Ac is a sialic acid side chain that has been shown to trigger the chemoreceptors that control sea anemone nematocyte firing (Ozacmak et al. 2001, Abdullah & Saad 2015). Sialic acids are nine-carbon sugars that can be either O- or N-glycosylated at the cell surface of mucins (Visser et al. 2021). There is large variability in the structural diversity and biological function of sialic acid, however, analysis of these glycans is challenging as there is very little known about their biosynthesis and function (Visser et al. 2021). Marcionetti et al. (2019) used genomic analysis to identify key proteins in the anemonefish epidermis that were positively selected for during anemonefish evolution. Maricionetti et al. (2019) found 17 genes to have evolved under positive selection at the origin of anemonefish evolution. They found that versican core proteins that are thought to bind N-acetylated sugars were positively selected for and thus may be used to mask detection of anemonefish by sea anemone chemoreceptors. The gene for the protein, O-GlcNAse, was also found to be positively selected and has the potential to cleave the sialic acid side chain creating neutral glycoproteins (as seen by Abdullah and Saad (2015)). The evolution of these proteins in anemonefish may have enhanced the establishment and maintenance of symbiosis between host sea anemones and anemonefish. Our study supports the findings of Marcionetti et al. (2019), that anemonefish have evolved mechanisms, such as alteration of glycans and cleaving of sugars, in order to reduce nematocyst discharge.

Anemonefish can generally acclimate to their host sea anemone in less than 24-48 hours (Mariscal 1970, Balamurugan et al. 2015, pers obv), and this is the point at which we would expect to see changes in the anemonefish mucus composition. However, this was not found in this study, rather it took three weeks of association with *E. quadricolor* before clear changes in the glycan composition of the anemonefish mucus were observed. Balamurugan et al. (2015) found that anemonefish secrete an intracellular mucous lining in the hypodermal region, that is not seen in *Terapon jarbua* (a fish species that does not associate with sea anemones). Therefore, perhaps it is not the external mucus layer that is the key to the initial formation of symbiosis, but rather that the internal mucus acts as a barrier to sea anemone nematocysts until the external mucus layer has the opportunity to adapt to the new environment through glycan changes, as we observed in this study. Mariscal (1970) found that after 20 hours isolated from a host sea anemone, anemonefish were stung (n=21) upon reintroduction to the host. We found that within 24 hours of host sea anemone removal, the glycan profile of the anemonefish mucus was no longer significantly different to the control. The loss of these changes in the glycan profile within 24 hours of removal from a host sea anemone, provides further support that anemonefish are not innately protected from sea anemone venom as previously thought (Hoepner et al. 2022) but indeed adjust their glycan profile when in association with a sea anemone host.

While the significant alteration of the glycan profile of anemonefish mucus is aligned with the significant decrease in nematocyte firing at anemonefish in association with the sea anemone host (after three weeks of association), we did not assess additional time points in the nematocyte experiment. In future studies, it would be important to assess the number of nematocytes fired at anemonefish mucus across shorter periods of 48 hours, one and two weeks. More comprehensive sampling during the acclimation period may shed light on the mechanisms by which hot sea anemones and anemonefishes adjust the nematocyst firing rate, and the chemical cues that initiate adaptive responses.

## 6. Conclusion

This study has provided a new piece in the puzzle of sea anemone and anemonefish symbiosis. Mucus from anemonefish not in association with sea anemones trigger high numbers of nematocytes from host sea anemones. Once living within association with a host sea anemone, anemonefish trigger significantly fewer nematocysts from host sea anemones than unacclimated anemonefish, which also corresponded with a significant change in the anemonefish glycan profile after three weeks of association. Triggering fewer nematocysts through glycan profile alterations, may be an important adaptation that has enabled anemonefish to live long-term in a host sea anemone, but due to the delay in glycan adaptation, another unknown component may contribute to the initial mechanism used by anemonefish to first enter their host sea anemone without being killed. Future studies are needed to identify the glycans in the anemonefish mucus that change during association with a host sea anemone and to investigate if the sialic acid content of acclimated anemonefish mucus is quantitatively lower than mucus of anemonefish not in association of a sea anemone. Thus, the chemistry behind the initial mechanism of entry for anemonefishes into a venomous environment still remains elusive.

## Acknowledgements

We would like to thank Leslie Morrison, Sarah Churcher and the Flinders University Biology Animal House staff for their support and expertise in sea anemone and anemonefish husbandry throughout the experiment. We would like to acknowledge the facilities, and the scientific and technical assistance of Microscopy Australia under the National Collaborative Research Infrastructure Strategy, at the South Australian Regional Facility, Flinders Microscopy and Microanalysis (FMMA), Flinders University. We would also like to thank Alex Sibley from Flinders Microscopy and Microanalysis for his expertise and assistance with Scanning Electron Microscopy and image analysis. The lipidomic and glycomic work was performed in part at the Melbourne Centre for Nanofabrication (MCN) in the Victorian Node of the Australian National Fabrication Facility (ANFF) we would like to acknowledge their scientific and technical assistance and facilities.

## Declarations

This research was supported by the Holsworth Endowment Fund and a Fisheries Society of the British Isles Small Research Grant awarded to CMH. The authors have no conflicts of interests to declare.

## Supplementary Materials

**Figure S1:**
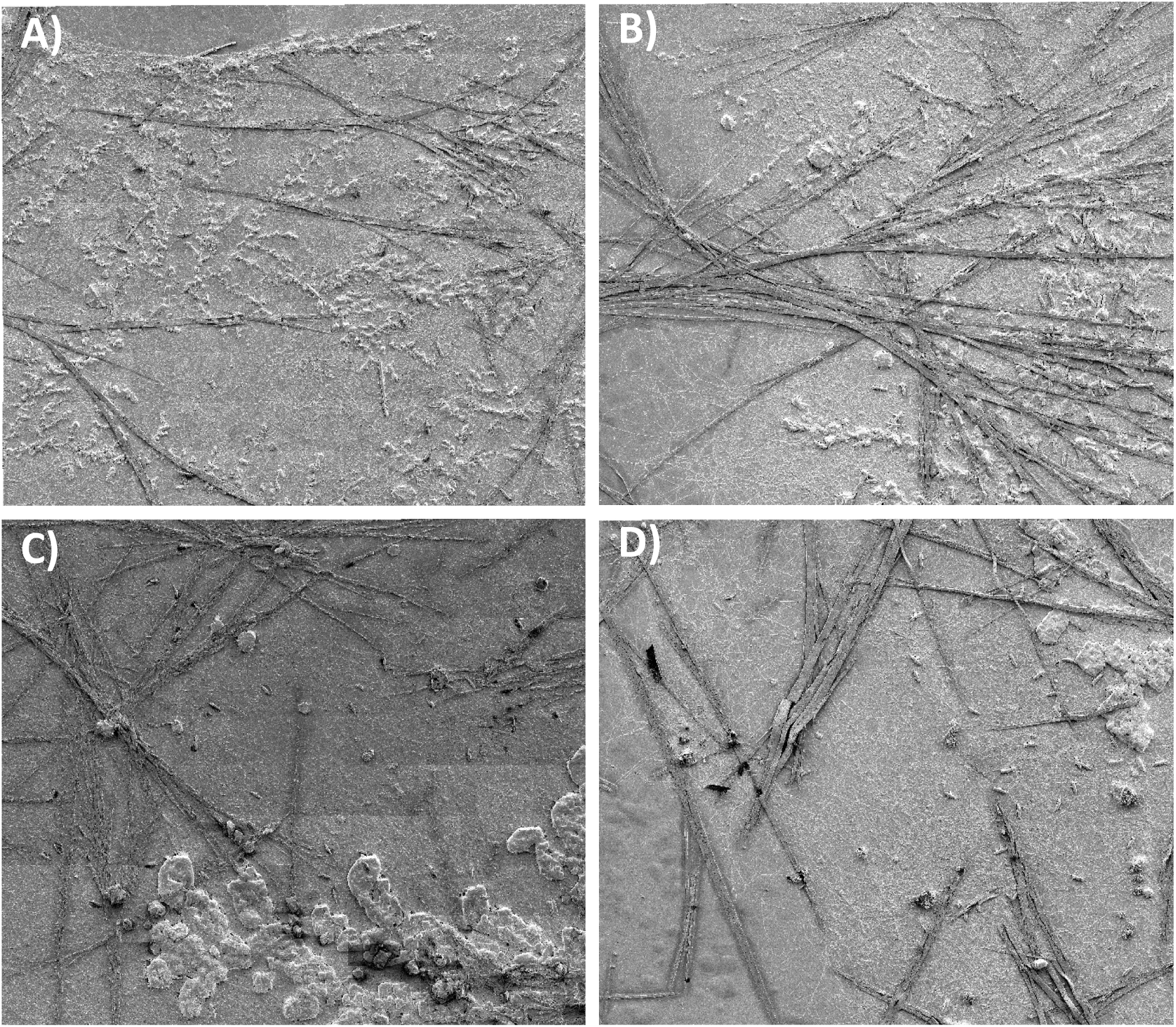
100 SEM images stitched together to represent 1mm2 from each glass slide A) prawn mucus before timepoint, B) damselfish mucus before timepoint, C) anemonefish mucus before association, D) anemonefish mucus with association

**Figure S2:**
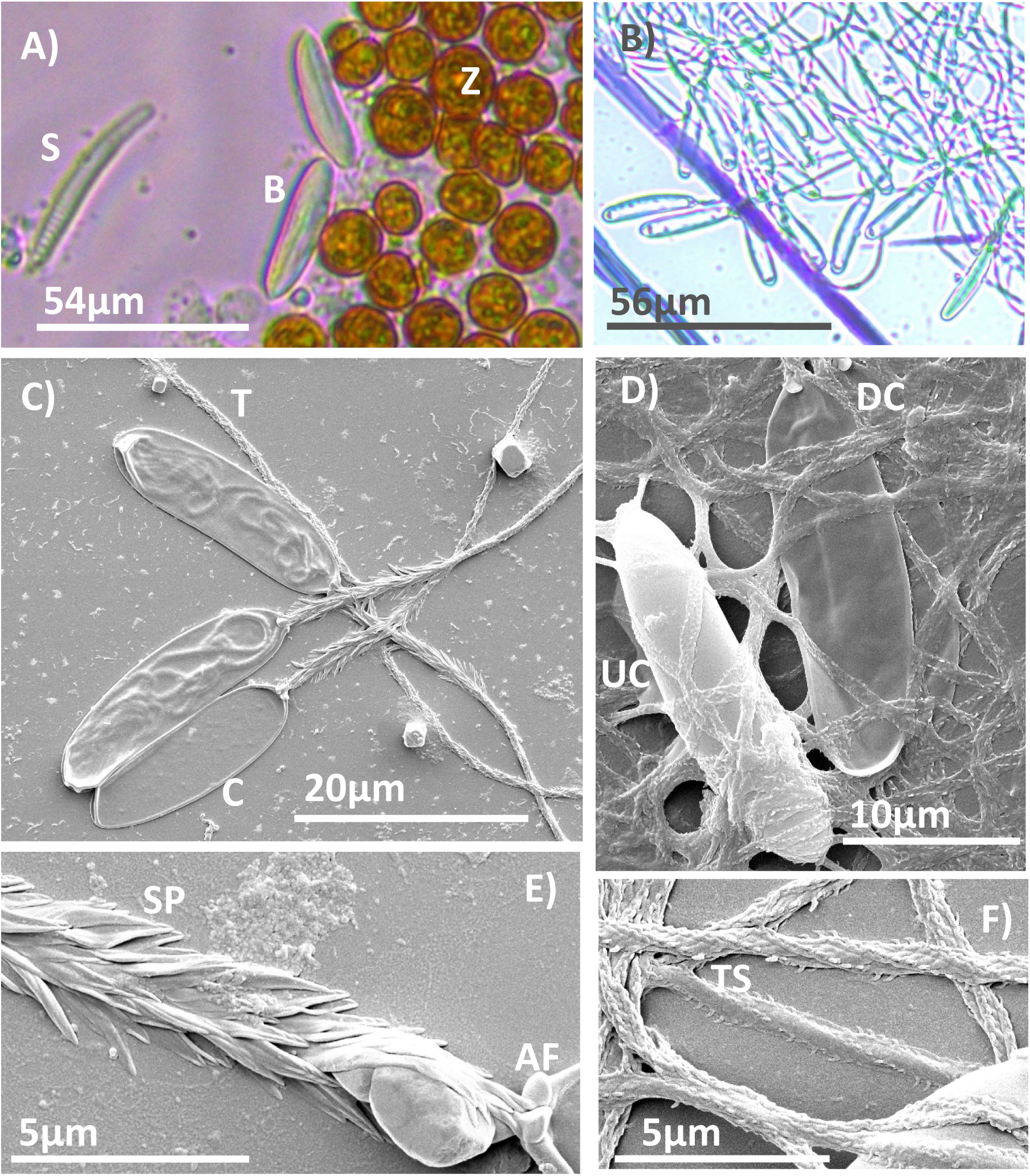
Light Microscopy and Scanning Electron Microscopy of basitrichous nematocytes from Entacmaea quadricolor. Image A) shows Light Microscopy of unfired nematocytes (basitrichous), spirocysts and zooxanthellae inside the tentacle. Image B) shows Light Microscopy of fired nematocytes (basitrichous) at 400x magnification. Image (C) shows SEM of fired basitrichous. Image D) shows SEM of fired capsules discharged and undischarged. Image E) shows SEM of the apical flaps which are open as the capsule is fired and spines at the base of the capsule. Image F) shows SEM of tubule spines which aid in anchoring to tissue.

**Figure S3:**
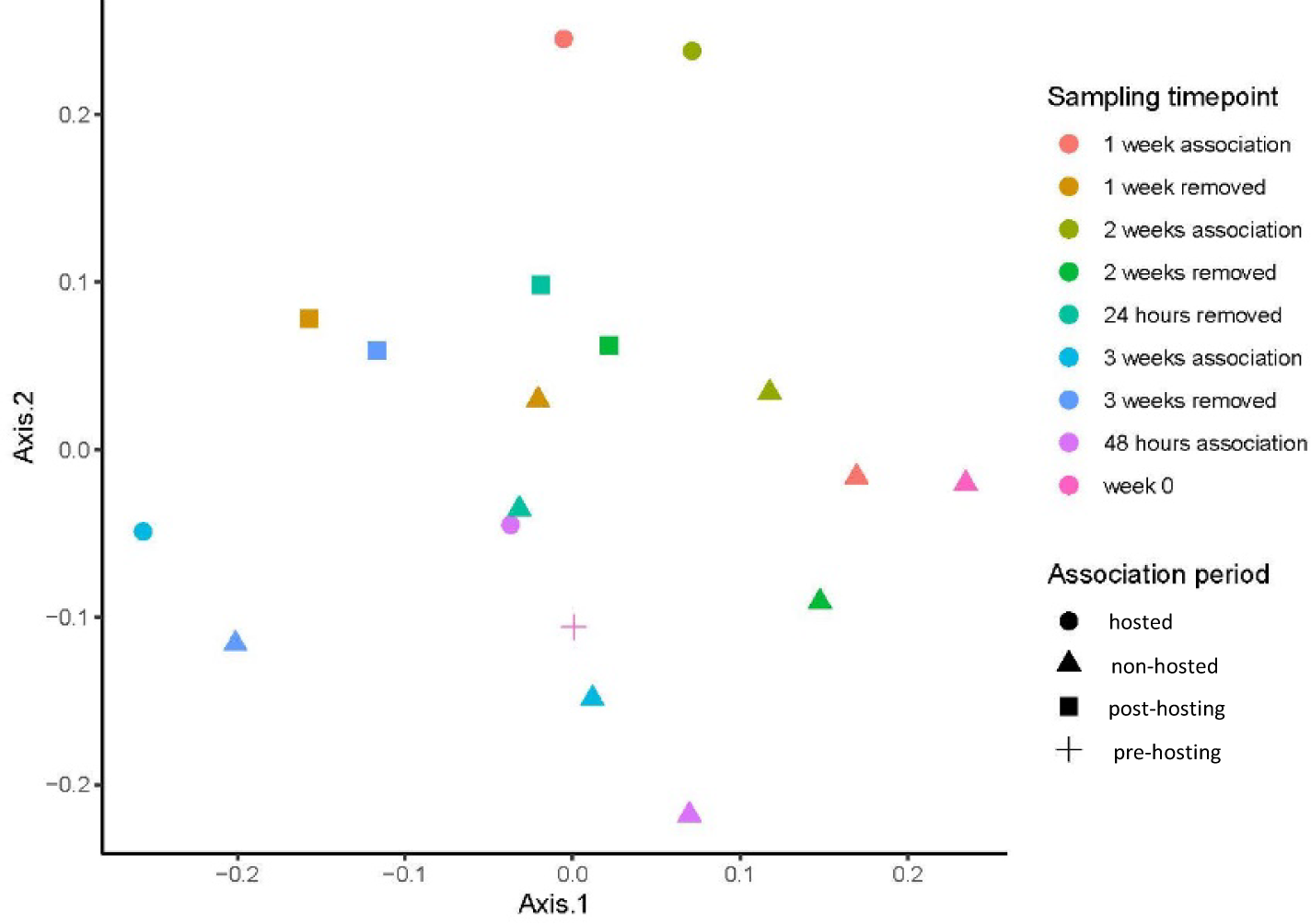
PCoA of *Amphiprion percula* mucus lipid profile across hosted and non-hosted periods with *E. quadricolor*

## Notes

### Competing Interest Statement

The authors have declared no competing interest.

